# Anti-Buoyancy Trap for Long-Term Quantitative Density Profiling of Adipocyte Spheroids

**DOI:** 10.64898/2026.03.10.708771

**Authors:** Min Young Kim, Sohae Yang, Minseok S. Kim

**Author notes:** Corresponding author*. *E-mail address:* (M. S. Kim).

## Abstract

3T3-L1 adipocyte spheroids are widely used three-dimensional adipose models that undergo pronounced physical property changes during adipogenic maturation, including progressive buoyancy driven by lipid accumulation. We quantitatively profiled the time-resolved evolution of bulk spheroid density over 60 days, revealing a 6.7% reduction from 1.022 to 0.954 g/cm³ associated with intracellular lipid accumulation. This buoyancy-driven displacement hinders standardized interrogation of spheroid physical properties during long-term culture. To enable longitudinal and reproducible analysis of density remodeling, we introduced an anti-buoyancy spheroid trap (AS-Trap) that mechanically stabilizes spheroids during extended culture. Morphometric analysis revealed that lipid droplet areas in mature spheroids (790.68±513.84 μm²) closely approximated in vivo adipocytes (846.34±257.28 μm²) and were substantially larger than those in 2D cultures, providing structural context for the observed density reduction. Notably, the terminal spheroid density converged toward reported values for native white adipose tissue, underscoring the physiological relevance of the material state achieved during maturation. Together, this work establishes density as a quantitative physical phenotype of adipogenic maturation and enables standardized long-term interrogation of adipose spheroids as living materials.

## Introduction

The global obesity epidemic drives urgent demand for physiologically relevant in vitro models to study adipose pathophysiology and develop therapeutic interventions [1–3]. Traditional two-dimensional (2D) adipocyte cultures, while foundational for understanding adipogenesis, fail to recapitulate the complex three-dimensional architecture, cell-cell interactions, and mechanical forces present in native adipose tissue [4–6]. These limitations include artificial substrate stiffness, lack of proper intercellular contacts, failure to develop characteristic unilocular morphology, and poor modeling of the heterogeneous cellular composition found in native adipose tissue [7, 8].

Three-dimensional adipocyte spheroids have emerged as superior alternatives that demonstrate enhanced adipogenic differentiation efficiency, improved insulin sensitivity, physiological hormone secretion profiles, and gene expression patterns closely resembling primary adipose tissue [9, 10]. Recent advances have shown that 3D adipocyte cultures better recapitulate metabolic functions and obesity-related pathophysiology compared to conventional 2D systems [11, 12]. Spheroids better model obesity-related inflammation and browning responses, providing platforms for evaluating metabolic therapies [13, 14].

However, a fundamental technical challenge has limited their widespread adoption: progressive density reduction during adipocyte maturation causes spheroids to become buoyant and float in culture media [15, 16]. This phenomenon creates critical experimental complications that severely limit spheroid utility for rigorous research applications [17, 18]. Inconsistent positioning disrupts automated imaging workflows essential for high-throughput screening, as floating spheroids move beyond the microscope’s working distance and focal plane [19]. Buoyant spheroids experience altered oxygen and nutrient gradients compared to settled cultures, with surface-floating spheroids receiving enhanced oxygenation while experiencing reduced access to nutrients that settle gravitationally [20]. Medium exchange procedures risk catastrophic spheroid loss, as aspiration and addition of fresh medium can inadvertently remove floating spheroids or damage them through turbulence [21]. Individual spheroid tracking becomes impossible when spheroids occupy unpredictable vertical positions within the medium column, preventing longitudinal studies of adipocyte maturation dynamics and dose-response relationships [22, 23]. Standardization across wells becomes impossible when spheroids occupy different vertical positions within the medium column, compromising experimental reproducibility and statistical power [24].

Previous attempts to address spheroid floating have employed various suboptimal solutions that compromise experimental integrity [25, 26]. Methylcellulose addition increases medium viscosity to prevent floating but fundamentally alters diffusion kinetics and may affect cellular metabolism through changes in nutrient and oxygen availability [27]. Matrigel embedding provides physical support but introduces undefined biological factors, significant batch-to-batch variability in composition, and potential confounding effects on adipocyte differentiation pathways [28]. Manual repositioning of spheroids using pipettes or other tools disrupts cultures, introduces operator-dependent variability, and proves impractical for large-scale studies requiring hundreds of spheroids [29]. Alternative approaches such as increasing seeding density or using specialized low-attachment surfaces fail to address the fundamental density-driven buoyancy while potentially altering spheroid formation kinetics [30]. These existing methods represent band-aid solutions that either alter the physiological culture environment or introduce uncontrolled variables that compromise data interpretation. Additionally, most approaches become increasingly ineffective as spheroids mature and achieve maximum buoyancy, limiting their utility for long-term studies. Critically, no studies have quantitatively characterized temporal density dynamics during spheroid maturation or developed engineering solutions maintaining physiological culture conditions.

Here, we present the first comprehensive quantitative analysis of density evolution in 3T3-L1 adipocyte spheroids over 60 days, revealing progressive density reduction from 1.022 to 0.954 g/cm³. We developed an innovative anti-buoyancy spheroid trap (AS-Trap) that prevents floating through mechanical retention (Figure 1A-B). Our mature spheroids achieve near-physiological density values and demonstrate superior lipid droplet morphology compared to conventional 2D cultures (Figure 1C), with mean droplet areas (790.68±513.84 μm²) closely approximating in vivo adipocyte dimensions (846.34±257.28 μm²) and significantly exceeding 2D culture values (376.09±196.18 μm²) (Figure 1D). This morphometric validation, coupled with the successful recapitulation of unilocular lipid storage characteristic of mature white adipocytes, establishes our spheroid system as a biomimetic model suitable for long-term metabolic research and drug discovery applications.

**Figure 1.**
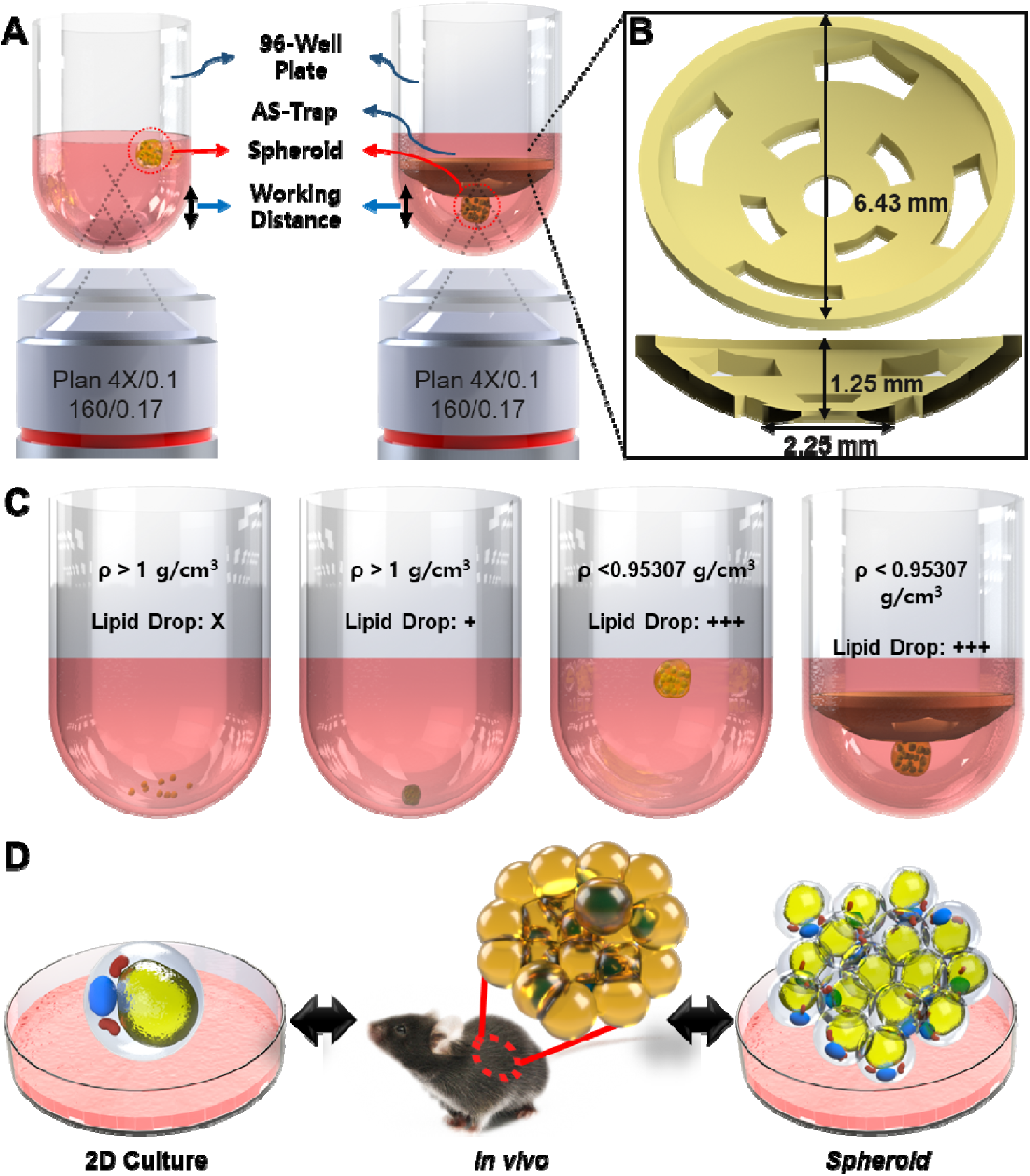
AS-Trap device for imaging buoyant 3T3-L1 adipocyte spheroids and conceptual comparison of culture models. (A) Schematic illustrating the challenge of imaging buoyant spheroids in a 96-well plate due to their tendency to float out of the microscope’s working distance (left). The AS-Trap device confines the spheroid within the working distance, enabling stable imaging (right). (B) Design and dimensions of the AS-Trap device. (C) Flotation occurs when ρ_spheroid < ρ_medium (ρ_medium ≈ 1.00–1.01 g/cm³ under typical culture conditions). (D) Comparison of cell morphology and lipid accumulation in conventional 2D culture, in vivo adipose tissue (mouse schematic), and 3D 3T3-L1 adipocyte spheroids, highlighting the 3D spheroid model’s aim to better mimic in vivo characteristics.

## Materials and Methods

### 3T3-L1 Cell Culture and Maintenance

3T3-L1 preadipocytes were purchased from the American Type Culture Collection (ATCC CL-173) and cultured according to standard protocols. Cells were grown in 96-well culture plates using pre-adipocyte culture medium consisting of Dulbecco’s Modified Eagle Medium (DMEM, Gibco) supplemented with 10% bovine calf serum (ATCC) and 1% Antibiotic-antimycotic solution 100× (HyClone) at 37°C in a humidified atmosphere with 5% CO . For adipogenic differentiation, cells were first allowed to proliferate for two days in pre-adipocyte medium. The medium was then replaced with adipocyte differentiation medium containing DMEM supplemented with antibiotics, 10% fetal bovine serum (Thermo Fisher), 1 μM dexamethasone (G-BIOSCIENCES), 0.5 mM 3-Isobutyl-1-methylxanthine (Sigma), and 1 μg/mL insulin (Sigma) for two days. Subsequently, cells were maintained in adipocyte maintenance medium (DMEM containing 10% FBS with 0.1 μg/mL insulin) for the duration of the experiment. For spheroid formation, 3T3-L1 preadipocytes were seeded at 5×10³ cells per well in ultra-low attachment 96-well round-bottom plates (Corning #7007) in 200 μL pre-adipocyte medium. Plates were centrifuged at 1200×g for 3 minutes to initiate aggregation, then incubated at 37°C, 5% CO . After 48 hours of spheroid formation, the same differentiation protocol was applied to spheroids as described for 2D cultures.

### Density Measurement by Sequential Vial

Spheroid density was determined using a sequential vial flotation method adapted from polymer characterization techniques (Figure 3A). Solutions of known density (0.85, 0.90, 0.95, 0.9524, 0.9526, 0.9531, 0.9535, 1.00, 1.05, 1.10 g/cm³) were prepared by mixing absolute ethanol and glycerol with distilled water at calculated ratios, with densities verified using a precision density meter (Density2Go, Mettler toledo). Individual density solutions (2 mL each) were prepared in separate glass vials and allowed to equilibrate at 26°C for 30 minutes before use. Individual spheroids were sequentially transferred through the density solutions, starting from the lowest density (0.85 g/cm³) and progressing to higher densities. Each spheroid was gently introduced into a vial using a wide-bore pipette tip and observed for 5 minutes to determine flotation behavior. The spheroid density was determined by identifying the critical density solution where the spheroid neither sank nor floated, remaining neutrally buoyant or at the threshold between sinking and floating. If a spheroid sank in one density solution but floated in the next lower density, the spheroid density was interpolated between these two values.

### AS-Trap Device Design and Fabrication

The anti-buoyancy spheroid trap (AS-Trap) was designed using computer-aided design software (SolidWorks 2023). Design criteria included: (1) secure spheroid retention up to 2 mm diameter, (2) compatibility with 96-well plate geometry, and (3) biocompatibility for long-term culture. The optimized design features a circular base (6.43 mm outer diameter) with curved retention arms arranged radially. Perfusion holes (2.25 mm width) between arms ensure adequate medium flow while maintaining spheroid retention. The device height (3.5 mm total) maintains spheroids within the working distance of standard objectives. Devices were fabricated using stereolithography (Form 3B, Formlabs) with HighTemp resin (Formlabs). Post-processing included washing in isopropyl alcohol (10 minutes), UV curing (60 minutes at 60°C), and sterilization by autoclaving (121°C, 20 minutes). Prior to use, AS-Trap devices were coated with Anti-adherent rinsing solution (StemCell Technologies) to prevent spheroid adhesion to the device surface. The devices were incubated with the anti-adherent solution for 1 hour at room temperature, then rinsed with sterile PBS before spheroid placement. This coating ensured that spheroids remained in suspension within the trap while being physically retained by the device architecture.

### Lipid Spot Staining and Lipid Quantification

Intracellular lipid droplets were visualized using LipidSpot™ (Biotium), which stains lipid droplets in live cells. For both 2D cultures and spheroids, LipidSpot™ and Hoechst (Invitrogen) were added to cells at a final concentration of 1× in adipocyte maintenance medium and incubated at 37°C for 30 minutes. For in vivo comparison, epididymal white adipose tissue was harvested from 12-week-old male C57BL/6J mice (n=5) maintained on standard chow diet. All animal procedures were approved by the Institutional Animal Care and Use Committee (DGIST-IACUC-23112802-0001). eWAT was isolated and stained with LipidSpot™ for 30 minutes. Following incubation, stained samples were washed three times with 1× DPBS (Welgene) to remove excess dye. Imaging was conducted using a Nikon A1R laser scanning confocal microscope (Nikon) with appropriate filter sets for LipidSpot™ fluorescence detection. Quantitative analysis of lipid droplet area was performed using IMARIS software (Bitplane) with the surface module to obtain detailed statistics for individual lipid droplets. Total lipid content per spheroid was calculated by summing all detected lipid droplet areas within each spheroid volume. For 2D cultures, lipid droplet area per cell was measured and averaged across multiple fields of view.

### Statistical Analysis

All data are presented as mean ± standard deviation (SD). Statistical analyses were performed using GraphPad Prism 9.0 (GraphPad Software, CA, USA). Comparisons between multiple groups were analyzed using one-way ANOVA followed by Tukey’s post-hoc test for multiple comparisons. Statistical significance was set at p < 0.05. All experiments were performed in triplicate with at least three independent biological replicates.

## Results

### Comparative Density Analysis Across Different Media Components

To define the physical boundary conditions governing spheroid buoyancy, we quantified the density of culture media components and reference solutions (Figure 3B). Absolute ethanol, used as a low-density reference, showed consistent measurements of 0.777±0.004 g/cm³ (n=5), providing the lower boundary for the density gradient columns. This value closely matches literature values for ethanol density at room temperature, validating the accuracy of our measurement system. Distilled water, serving as a near-neutral reference, measured 0.993±0.002 g/cm³, slightly below the theoretical value of 1.000 g/cm³ due to temperature effects and the precision of the measurement system. Glycerol, used as a high-density reference, exhibited measurements of 1.272±0.002 g/cm³, providing the upper boundary for density comparisons. Standard DMEM medium, which forms the base for all culture media used in the study, showed a density of 1.000±0.002 g/cm³. This value defines the density boundary separating sinking and floating regimes for spheroids in standard culture conditions. Adipocyte maintenance medium, which contains additional components including serum proteins, growth factors, and hormones, demonstrated a slightly higher density of 1.01±0.002 g/cm³. These medium density measurements are particularly important for predicting spheroid behavior during different phases of the culture protocol. During medium changes, the exposure to fresh maintenance medium (1.01 g/cm³) means that spheroids densities below approximately 1.000 g/cm³ before consistent floating is observed. This explains the observation that spheroid floating becomes problematic around weeks 4 to 5, when densities reach this critical threshold. Consequently, these measurements establish a quantitative physical framework linking spheroid density evolution to medium-dependent buoyant behavior.

### AS-Trap Device Implementation and Spheroid Retention

The implementation of the AS-Trap device successfully addressed the critical technical challenge of spheroid floating while maintaining the physiological characteristics that make spheroids valuable as adipose tissue models. Throughout the 60-day culture period, AS-Trap devices demonstrated consistent performance in retaining spheroids at predictable positions within culture wells, enabling standardized imaging and experimental procedures. Physical retention of spheroids was achieved without observable adverse effects on morphological development. Spheroids cultured within AS-Trap devices followed identical growth kinetics to free-floating controls, reaching comparable final diameters and maintaining the characteristic translucent appearance of mature adipocytes. The open architecture of the device, with its 2.25 mm perfusion channels, permitted unimpeded medium exchange during routine feeding procedures, ensuring that retained spheroids received adequate nutrient supply and waste removal.

Visual assessment during medium changes confirmed complete medium replacement within the device structure, with no observable dead zones or areas of poor perfusion. The curved retention arms successfully accommodated spheroids throughout their growth from initial around 1,000 μm diameter aggregates to final sizes exceeding over 1,500 μm, demonstrating the scalability of the device design across the full range of spheroid development. The anti-adherent coating applied to device surfaces proved effective in preventing spheroid attachment to the plastic surfaces, ensuring that spheroids remain in suspension while being physically contained within the device boundaries. This maintained the three-dimensional culture characteristics essential for proper adipocyte development while solving the practical challenges associated with spheroid floating and loss during medium changes.

## Temporal Evolution of Spheroid Morphology, Size, and Mass

3T3-L1 adipocyte spheroids demonstrated remarkable morphological transformation over the 60-day culture period, providing insights into long-term adipocyte development in three-dimensional culture. Initial aggregates formed within 48 hours of seeding appeared as compact, dark spheres with uniform cellular density and well-defined boundaries (Figure 2A). During the first phase of culture (Days 1-4), corresponding to the growth period before differentiation induction, spheroids showed modest size increases from 409.37±9.83 μm to 481.62±21.83 μm (Figure 4A). This initial growth reflected continued cell proliferation within the three-dimensional structure, with spheroids maintaining their compact, opaque appearance characteristic of undifferentiated preadipocytes. Following adipogenic induction on Day 4, spheroids entered a distinct differentiation phase (Days 4-18) characterized by progressive morphological changes and size expansion. By Day 11, spheroids had reached 721.56±17.04 μm in diameter, representing a significant acceleration in growth rate compared to the pre-differentiation period. This expansion continued through Day 18, reaching 800.96±20.21 μm. During this phase, spheroids began developing the characteristic translucent appearance indicative of early lipid accumulation, with the peripheral regions becoming increasingly transparent while maintaining defined spherical morphology (Figure 2A-D). The most dramatic changes occurred during the maturation phase (Days 18-46), where spheroids underwent accelerated growth coupled with pronounced morphological transformation. Diameter measurements revealed a steep increase from 800.96±20.21 μm at Day 18 to

**Figure 2.**
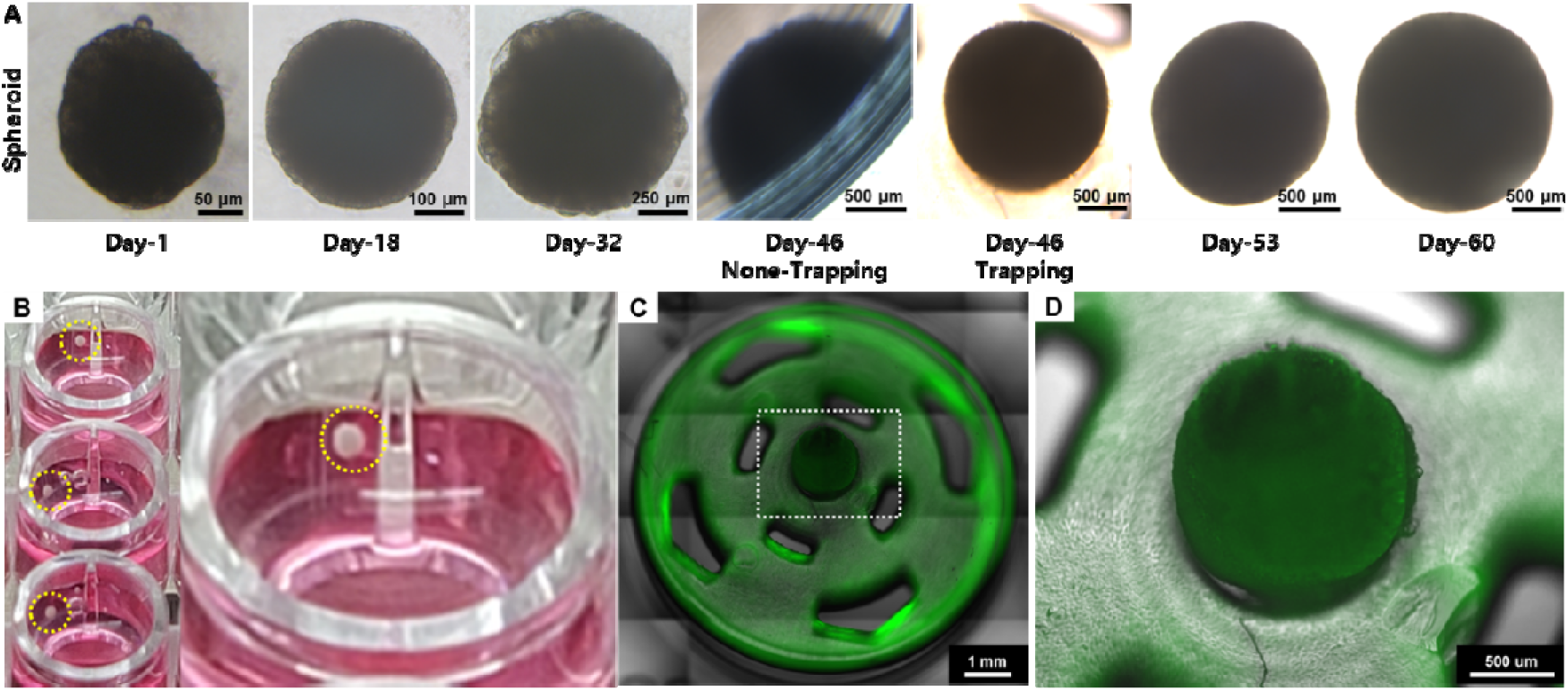
Morphological evolution of 3T3-L1 adipocyte spheroids and functionality of the AS-Trap. (A) Time-course bright-field microscopy images of 3T3-L1 adipocyte spheroids showing morphological changes and growth from Day 1 to Day 60. (B) Photographs showing the 96-well plate containing 3T3-L1 spheroids (indicated by yellow dashed circles) in culture medium. (C) Fluorescence image of a LipidSpot^TM^ (green, for lipids) 3T3-L1 spheroid captured within the AS-Trap. (D) Magnified view of spheroid trapped by the AS-Trap.

**Figure 3.**
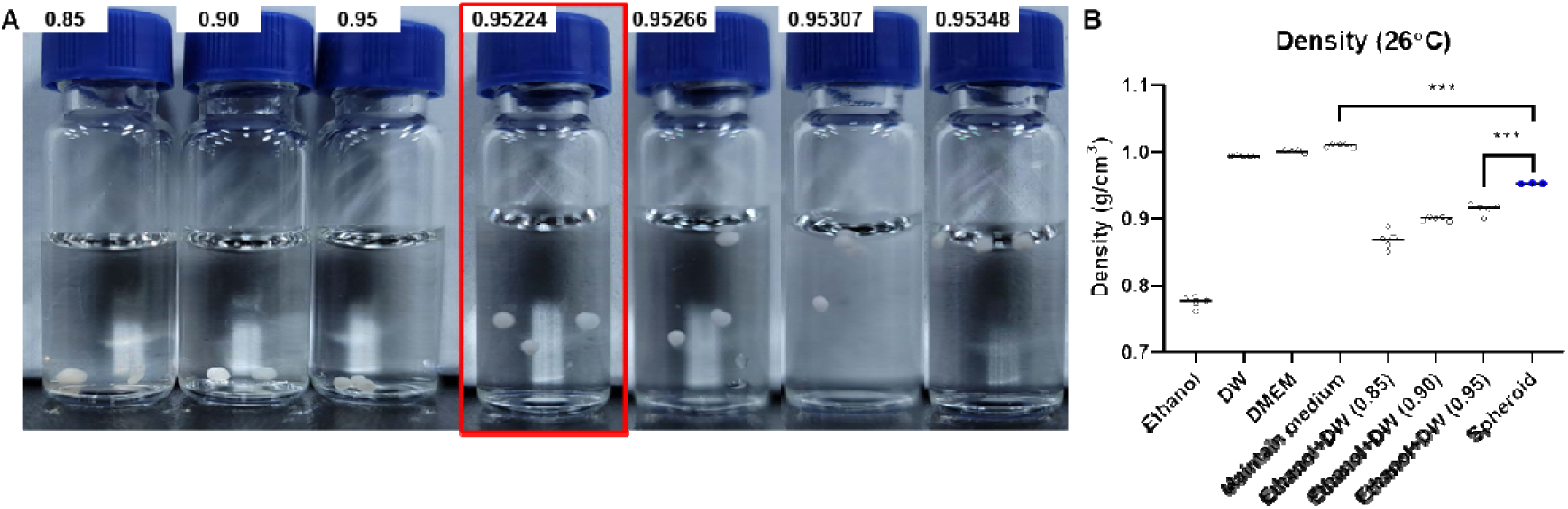
Density determination of 3T3-L1 adipocyte spheroids. (A) Visual assessment of 3T3-L1 spheroid buoyancy in ethanol-water solutions of varying densities (g/cm³ indicated above vials: 0.85, 0.90, 0.95, 0.95224, 0.95266, 0.95307, 0.95348). Spheroids are suspended in the solution with a density of 0.95224 g/cm³ (highlighted by red box), indicating their density is close to this value. (B) Quantitative comparison of spheroid density with densities of various reference solutions at 26°C. Data points represent individual measurements. Bars indicate mean ± SEM. Statistical significance indicated: ***p < 0.001 (One-way ANOVA with Tukey’s post-hoc test comparing Spheroid group to all other groups).

**Figure 4.**
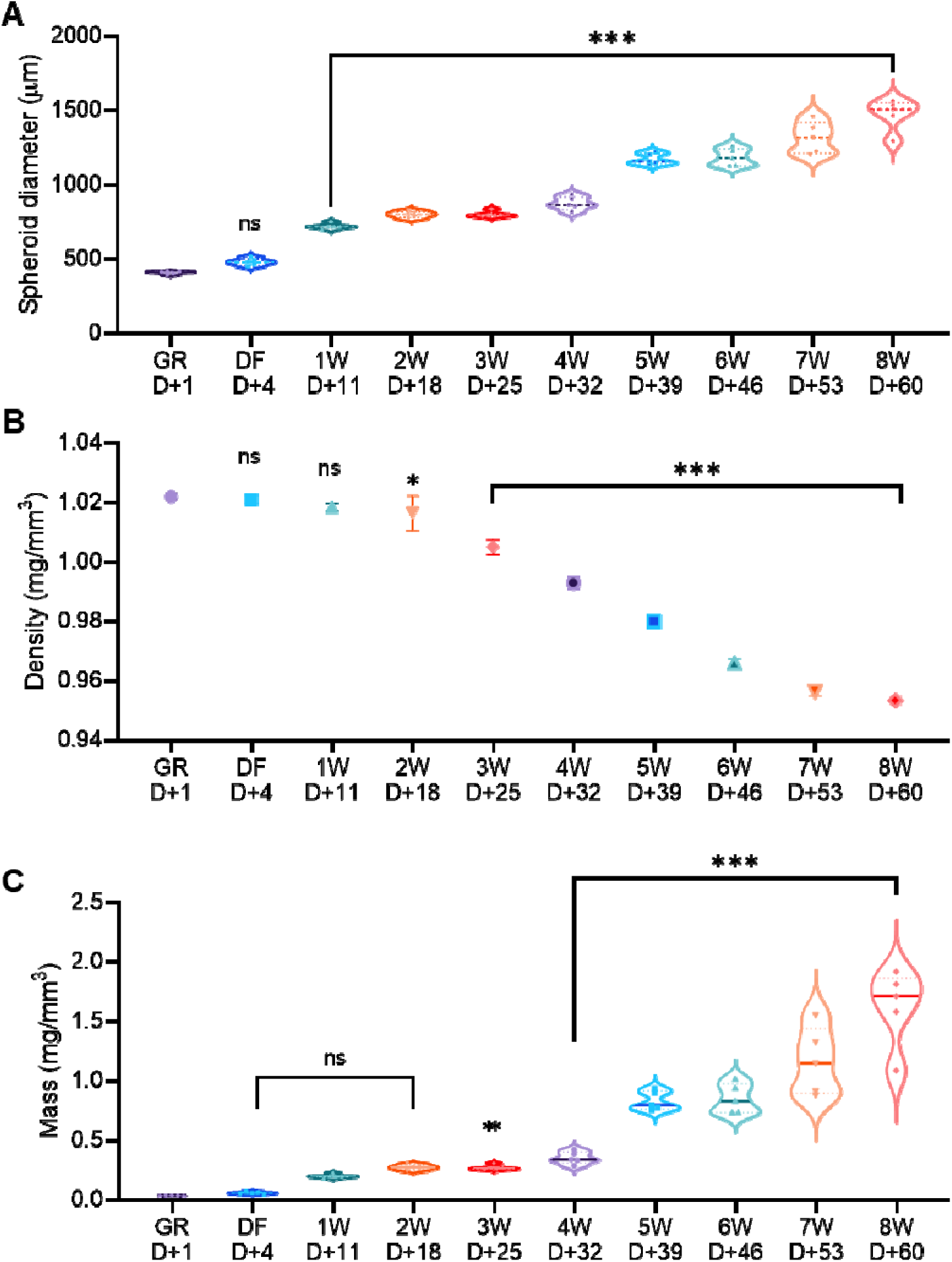
Temporal changes in biophysical properties of 3T3-L1 adipocyte spheroids during 60 days of culture. (A) Spheroid diameter over time. Violin plots show the distribution of spheroid diameters at indicated time points: GR D+1 (Growth Day 1), DF D+4 (Differentiation Day 4), and weekly from 1W D+11 to 8W D+60. (B) Spheroid density over time. Individual data points are shown for each time point. A significant decrease in density is observed, particularly after week 2. (C) Spheroid mass over time. Violin plots show the distribution of spheroid mass at indicated time points. Statistical significance indicated: ns (not significant), *p < 0.05, **p < 0.01, and ***p < 0.001 (One-way ANOVA with Tukey’s post-hoc test).

1,186.87±52.08 μm by Day 46, representing a 32.5% increase over this 28-day period. Growth continued beyond Day 46, though at a reduced rate, reaching a final diameter of 1,475.07±95.64 μm at Day 60. This represents an overall 3.6-fold increase in diameter from initial aggregation, corresponding to an approximately 47-fold increase in spheroid volume assuming spherical geometry. In the later stages (days 46-60), there was some variability between individual spheroids and no further increase in size. These results suggest that spheroids had reached near-maximum sustainable size for the given culture conditions.

Mass provided complementary insights into spheroid development, revealing the dramatic accumulation of cellular material over the culture period (Figure 4C). Initial spheroid mass of 0.037±0.002 mg reflected the compact cellular aggregates with minimal extracellular space. During the differentiation phase, mass increased steadily to 0.274±0.019 mg by Day 18, representing a 7.45-fold increase that accompanied the morphological changes and early lipid accumulation. The maturation phase showed exponential mass accumulation, reaching 0.85±0.112 mg by Day 46. This 14-fold increase over the 42-day maturation period corresponded with the period of maximum lipid droplet formation and spheroid translucency development. The final mass measurement at Day 60 of 1.622±0.29 mg represented an overall 44-fold increase from initial values, demonstrating the remarkable capacity for lipid storage in the three-dimensional culture system. Interestingly, the mass-to-diameter relationship revealed important insights into spheroid density changes over time.

### Quantitative Density Profiling During Adipocyte Maturation

Direct measurements of bulk spheroid density using a graded density flotation method revealed pronounced biophysical remodeling during adipocyte maturation (Figure 4B). This analysis establishes density as a time-resolved physical phenotype of adipogenic maturation in three-dimensional adipocyte spheroids. Undifferentiated spheroids exhibited a uniform initial density of 1.022±0.001 g/cm³ (n=5), which was consistent across batches and experimental replicates. This value is consistent with hydrated cellular tissues dominated by aqueous and proteinaceous components. The consistency of these measurements, with standard errors below 0.1%, demonstrated the reliability of the density measurement technique and the uniformity of spheroid formation across wells. During the early growth phase (Days 1–4), spheroid density remained stable (1.022±0.001 g/cm³), indicating that volumetric expansion at this stage was dominated by cell proliferation rather than compositional remodeling. This stability continued through the early differentiation period, with Day 11 measurements showing 1.019±0.001 g/cm³ and Day 18 measurements of 1.019±0.001 g/cm³. The minimal density changes during this 17-day period, despite significant increases in spheroid size and the initiation of adipogenic differentiation, suggests that early differentiation events involve primarily transcriptional and protein expression changes without substantial lipid accumulation.

The onset of the maturation phase coincided with the first measurable reduction in spheroid density, reaching 1.005±0.002 g/cm³ by Day 25. This 1.6% reduction from baseline represented the onset of substantial intracellular lipid accumulation, as triglycerides began replacing aqueous cytoplasmic components. Density reduction accelerated between Days 25 and 39, crossing the 1.000 g/cm³ threshold by Day 32 and reaching 0.980±0.002 g/cm³ by Day 39. The 1.6% density reduction over this 7-day period represented the steepest rate of change observed throughout the entire culture period. The most pronounced density reduction occurred between Days 39-46, reaching 0.966±0.001 g/cm³. This regime corresponds to a transition in bulk material composition, consistent with extensive lipid accumulation replacing aqueous cytoplasmic volume. Spheroids with densities at or below this threshold showed a tendency to float to the medium surface, creating the technical challenges that motivated the development of the AS-Trap system. Beyond Day 46, density continued to decline but at a reduced rate, reaching 0.957±0.001 g/cm³ by Day 53 and 0.954±0.001 g/cm³ at the final Day 60 measurement. This plateau effect suggests that spheroids had reached a metabolic equilibrium where lipid synthesis and turnover had stabilized. The final density of 0.954 g/cm³ represents a total reduction of 6.7% from initial values, a remarkable change that reflects the extensive lipid accumulation characteristic of mature white adipocytes [31]. The consistency of density measurements across individual spheroids, as indicated by the small standard errors throughout the time course, demonstrated that the density reduction was a uniform characteristic of the adipogenic maturation process rather than variable behavior between individual spheroids. The low variance observed across the time course indicates that density remodeling is a reproducible and intrinsic feature of adipogenic maturation in spheroid culture.

### Lipid Droplet Morphology: Bridging 2D Culture and In Vivo Tissue

Detailed morphometric analysis of lipid droplets provided crucial validation of the physiological relevance of the spheroid model by comparing lipid storage characteristics across different culture systems and native tissue (Figure 5). This analysis revealed striking differences that highlight the advantages of three-dimensional culture for recapitulating in vivo adipocyte characteristics. In standard 2D 3T3-L1 cultures, adipocytes developed numerous small lipid droplets with a mean area of 376.09±196.18 μm² (Figure 5B). The size distribution in 2D cultures was relatively narrow, ranging from 99.6 to 943.1 μm², with most droplets clustering in the smaller size ranges. The multilocular appearance characteristic of 2D adipocyte cultures persisted throughout extended differentiation periods, with cells maintaining multiple small to medium-sized lipid droplets rather than progressing to the large single lipid droplets morphology typical of mature white adipocytes in vivo.

**Figure 5.**
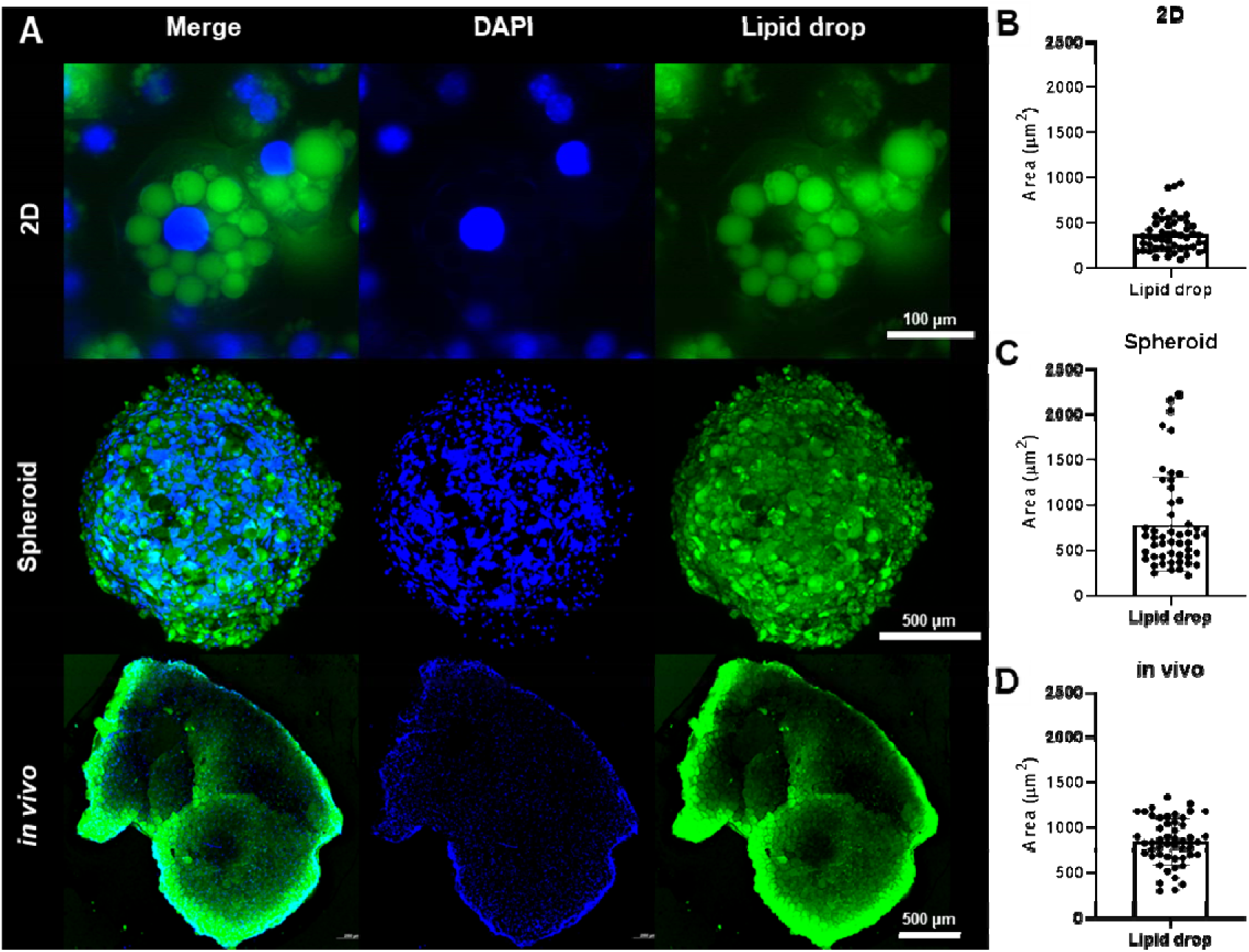
Comparative analysis of lipid droplet morphology in 2D cultured 3T3-L1 cells, 3T3-L1 spheroids, and in vivo murine adipose tissue. (A) Representative confocal images. 2D cultured 3T3-L1 adipocytes, Day 60 3T3-L1 adipocyte spheroid, and in vivo mouse white adipose tissue. DAPI (blue), and Lipid droplet (Lipid Spot^™^, green). (B) Quantification of lipid droplet area in 2D cultured 3T3-L1 adipocytes. (C) Quantification of lipid droplet area in Day 60 3T3-L1 adipocyte spheroids. (D) Quantification of lipid droplet area in vivo murine white adipose tissue.

Analysis of individual droplet characteristics in 2D cultures revealed several limitations of the monolayer system. The mean droplet area of 376.09 μm² represents less than half the size achieved in either spheroid culture or native tissue. Spheroid adipocytes displayed dramatically different lipid droplet characteristics, with significantly larger droplets averaging 790.68±513.84 μm² (Figure 5C). This represents a 2.8-fold increase in mean droplet area compared to 2D cultures, approaching the unilocular morphology characteristic of mature white adipocytes. The size distribution in spheroids was notably broader, ranging from 223.6 to 2,224.3 μm². This capacity for generating large, mature lipid droplets represents a significant advancement in adipocyte modeling, as the size and unilocular morphology of lipid droplets are key characteristics distinguishing mature adipocytes from preadipocytes or incompletely differentiated cells. For in vivo comparison, fresh epididymal white adipose tissue from healthy adult mice exhibited lipid droplets with a mean area of 846.34±257.28 μm² (Figure 5D). The size distribution ranged from 300.7 to 1,339.4 μm². Statistical analysis revealed no significant difference between spheroids and in vivo droplet areas (p=0.12), demonstrating that the spheroid culture system successfully recapitulates the lipid storage characteristics of native adipose tissue [32, 33]. The close correspondence between spheroid and in vivo lipid droplet sizes represents a crucial validation of the spheroid model’s physiological relevance. The ability to achieve in vivo-like lipid droplet morphology in vitro opens new possibilities for studying adipocyte biology, lipid metabolism, and therapeutic interventions in a controlled, reproducible system that maintains key characteristics of native tissue.

## Discussion

This study provides the first quantitative characterization of density evolution during long-term 3T3-L1 adipocyte spheroid culture and presents an engineering solution to overcome buoyancy-induced technical challenges. Our findings reveal that 3T3-L1 adipocyte spheroids undergo progressive density reduction from 1.022 to 0.954 g/cm³ over 60 days, representing a 6.7% decrease that correlates with intracellular lipid accumulation and buoyancy development[31]. The temporal density profile demonstrates distinct maturation phases consistent with known adipogenic differentiation pathways [34, 35]. During the initial 18 days, density remained stable (1.022 to 1.019 g/cm³) despite size increases from 409.37±9.83 μm to 800.96±20.21 μm, indicating early expansion was driven by cell proliferation rather than lipid accumulation [36, 37]. Measurable density reduction began around Day 25, with the most rapid decline occurring between Days 32-46, coinciding with spheroid translucency development. The critical threshold density of 1.000 g/cm³, below which spheroids exhibited buoyancy in culture medium (1.010±0.002 g/cm³), was crossed around Day 32, explaining why floating typically emerges during weeks 4-5 of culture [12]. Our comparative lipid droplet analysis validates the spheroid model’s physiological relevance [38, 39]. Mean lipid droplet area in spheroids (790.68±513.84 μm²) showed no statistical difference from in vivo adipose tissue (846.34±257.28 μm², p=0.12), while both were significantly larger than 2D cultures (376.09±196.18 μm²). This 2.1-fold increase compared to 2D cultures indicates successful recapitulation of unilocular morphology characteristic of mature white adipocytes [40, 41]. Recent studies have confirmed that 3D adipocyte models better preserve physiological characteristics and metabolic functions compared to conventional 2D systems [11, 12]. The AS-Trap device successfully addressed spheroid buoyancy while preserving physiological culture conditions. The open architecture with 2.25 mm perfusion channels maintained adequate mass transport, enabling spheroid retention throughout the 60-day culture period without compromising growth or differentiation. This compatibility with standard 96-well plates enables standardized imaging protocols and longitudinal studies previously impossible due to spheroid floating. The 44-fold mass increase over 60 days, coupled with density measurements showing standard errors below 0.1%, demonstrates the reproducibility of adipocyte maturation in spheroid culture. The final density of 0.953 g/cm³ approaches physiological white adipose tissue values, validating spheroids as biomimetic models.

Current challenges in 3D spheroid culture include inconsistent formation, variable sizes, and technical difficulties in long-term maintenance [42, 43]. Our AS-Trap system addresses these limitations by providing a standardized platform for spheroid culture that maintains physiological conditions while preventing buoyancy-related complications. This advancement is particularly relevant given the growing recognition of spheroids as essential tools for drug discovery and disease modeling [44, 45]. The implications of this work extend beyond technical improvements in spheroid culture methodology. By enabling reliable long-term culture of mature adipocyte spheroids, our platform facilitates studies of adipose tissue biology that were previously technically challenging or impossible [46, 47]. The quantitative density profiling approach developed here could be applied to other spheroid systems where density changes affect experimental outcomes [48]. Furthermore, the AS-Trap design principles could be adapted for different spheroid types and culture applications, providing a generalizable solution to buoyancy challenges in 3D culture systems [49].

## Conclusions

This study establishes fundamental biophysical parameters for 3T3-L1 adipocyte spheroid culture and provides practical solutions to technical barriers that have limited their adoption. By achieving near-physiological density (0.954 g/cm³) and lipid droplet morphology comparable to in vivo tissue (790.68±513.84 vs 846.34±257.28 μm²), our spheroid model bridges the gap between reductionist 2D culture and complex animal models. The AS-Trap platform enables standardized, long-term culture essential for mechanistic studies and drug discovery applications. These advances position adipocyte spheroids as powerful tools for investigating metabolic disease mechanisms and developing therapeutic interventions for the global obesity epidemic.

## CRediT Authorship Contribution Statement

**Min Young Kim:** Writing – review & editing, Writing – original draft, Visualization, Conceptualization, Methodology, Investigation, Data curation. **Minseok S. Kim:** Writing – review & editing, Writing – original draft, Supervision, Conceptualization, Project administration, Funding acquisition.

## Declaration of Competing Interest

The authors declare that they have no known competing financial interests or personal relationships that could have appeared to influence the work reported in this paper.

## Acknowledgements

This work was supported by the National Research Foundation (NRF) of Korea, funded by the Korean government (MSIP) (No. 2022R1A2C2091870).

## Data Availability

Data will be made available on request.

## Notes

### Competing Interest Statement

The authors have declared no competing interest.

## References

[1] A. Chait, L.J. Den Hartigh, Adipose Tissue Distribution, Inflammation and Its Metabolic Consequences, Including Diabetes and Cardiovascular Disease, Frontiers in Cardiovascular Medicine 7 (2020).

[2] M. Longo, F. Zatterale, J. Naderi, L. Parrillo, P. Formisano, G.A. Raciti, F. Beguinot, C. Miele, Adipose Tissue Dysfunction as Determinant of Obesity-Associated Metabolic Complications, International Journal of Molecular Sciences 20(9) (2019) 2358.

[3] G.D.a.I. Collaborators, Global burden of 369 diseases and injuries in 204 countries and territories, 1990–2019: a systematic analysis for the Global Burden of Disease Study 2019, The Lancet 396(10258) (2020) 1204–1222.

[4] Evan, Bruce, What We Talk About When We Talk About Fat, Cell 156(1-2) (2014) 20–44.

[5] A.L. Ghaben, P.E. Scherer, Adipogenesis and metabolic health, Nature Reviews Molecular Cell Biology 20(4) (2019) 242–258.

[6] A. Sakers, M.K. De Siqueira, P. Seale, C.J. Villanueva, Adipose-tissue plasticity in health and disease, Cell 185(3) (2022) 419–446.

[7] F. Ruiz-Ojeda, A. Rupérez, C. Gomez-Llorente, A. Gil, C. Aguilera, Cell Models and Their Application for Studying Adipogenic Differentiation in Relation to Obesity: A Review, International Journal of Molecular Sciences 17(7) (2016) 1040.

[8] S. Muller, I. Ader, J. Creff, H. Leménager, P. Achard, L. Casteilla, L. Sensebé, A. Carrière, F. Deschaseaux, Human adipose stromal-vascular fraction self-organizes to form vascularized adipose tissue in 3D cultures, Scientific Reports 9(1) (2019).

[9] P.A. Turner, B. Gurumurthy, J.L. Bailey, C.M. Elks, A.V. Janorkar, Adipogenic differentiation of human adipose-derived stem cells grown as spheroids, Process Biochemistry 59 (2017) 312–320.

[10] J. Dufau, J.X. Shen, M. Couchet, T. De Castro Barbosa, N. Mejhert, L. Massier, E. Griseti, E. Mouisel, E.-Z. Amri, V.M. Lauschke, M. Rydén, D. Langin, In vitro and ex vivo models of adipocytes, American Journal of Physiology-Cell Physiology 320(5) (2021) C822–C841.

[11] A. Ioannidou, S. Alatar, R. Schipper, F. Baganha, M. Åhlander, A. Hornell, R.M. Fisher, C.E. Hagberg, Hypertrophied human adipocyte spheroids as *in vitro* model of weight gain and adipose tissue dysfunction, The Journal of Physiology 600(4) (2022) 869–883.

[12] T.M. Avelino, M.G.-A. Provencio, L.A. Peroni, R.R. Domingues, F.R. Torres, P.S.L. De Oliveira, A.F.P. Leme, A.C.M. Figueira, Improving obesity research: Unveiling metabolic pathways through a 3D In vitro model of adipocytes using 3T3-L1 cells, PLOS ONE 19(5) (2024) e0303612.

[13] L.S. Baptista, K.R. Silva, L. Jobeili, L. Guillot, D. Sigaudo-Roussel, Unraveling White Adipose Tissue Heterogeneity and Obesity by Adipose Stem/Stromal Cell Biology and 3D Culture Models, Cells 12(12) (2023) 1583.

[14] K. Endo, T. Sato, A. Umetsu, M. Watanabe, F. Hikage, Y. Ida, H. Ohguro, M. Furuhashi, 3D culture induction of adipogenic differentiation in 3T3-L1 preadipocytes exhibits adipocyte-specific molecular expression patterns and metabolic functions, Heliyon 9(10) (2023) e20713.

[15] M. Vitacolonna, R. Bruch, A. Agaçi, E. Nürnberg, T. Cesetti, F. Keller, F. Padovani, S. Sauer, K.M. Schmoller, M. Reischl, M. Hafner, R. Rudolf, A multiparametric analysis including single-cell and subcellular feature assessment reveals differential behavior of spheroid cultures on distinct ultra-low attachment plate types, Frontiers in Bioengineering and Biotechnology 12 (2024).

[16] A.P.P. Guimaraes, I.R. Calori, H. Bi, A.C. Tedesco, SpheroMold: modernizing the hanging drop method for spheroid culture, Frontiers in Drug Delivery 4 (2024).

[17] O. Sirenko, T. Mitlo, J. Hesley, S. Luke, W. Owens, E.F. Cromwell, High-Content Assays for Characterizing the Viability and Morphology of 3D Cancer Spheroid Cultures, ASSAY and Drug Development Technologies 13(7) (2015) 402–414.

[18] T.H. Booij, L.S. Price, E.H.J. Danen, 3D Cell-Based Assays for Drug Screens: Challenges in Imaging, Image Analysis, and High-Content Analysis, SLAS Discovery 24(6) (2019) 615–627.

[19] J.S.K. Yuen Jr, A.J. Stout, N.S. Kawecki, S.M. Letcher, S.K. Theodossiou, J.M. Cohen, B.M. Barrick, M.K. Saad, N.R. Rubio, J.A. Pietropinto, H. Dicindio, S.W. Zhang, A.C. Rowat, D.L. Kaplan, Perspectives on scaling production of adipose tissue for food applications, Biomaterials 280 (2022) 121273.

[20] R.J. McMurtrey, Analytic Models of Oxygen and Nutrient Diffusion, Metabolism Dynamics, and Architecture Optimization in Three-Dimensional Tissue Constructs with Applications and Insights in Cerebral Organoids, Tissue Engineering Part C: Methods 22(3) (2016) 221–249.

[21] A.Y. Hsiao, Y.C. Tung, X. Qu, L.R. Patel, K.J. Pienta, S. Takayama, 384 hanging drop arrays give excellent *Z* factors and allow versatile formation of co culture spheroids, Biotechnology and Bioengineering 109(5) (2012) 1293–1304.

[22] R. Foty, A Simple Hanging Drop Cell Culture Protocol for Generation of 3D Spheroids, Journal of Visualized Experiments (51) (2011).

[23] Y.-C. Tung, A.Y. Hsiao, S.G. Allen, Y.-S. Torisawa, M. Ho, S. Takayama, High-throughput 3D spheroid culture and drug testing using a 384 hanging drop array, The Analyst 136(3) (2011) 473–478.

[24] K. Alessandri, B.R. Sarangi, V.V. Gurchenkov, B. Sinha, T.R. Kießling, L. Fetler, F. Rico, S. Scheuring, C. Lamaze, A. Simon, S. Geraldo, D. Vignjević, H. Doméjean, L. Rolland, A. Funfak, J. Bibette, N. Bremond, P. Nassoy, Cellular capsules as a tool for multicellular spheroid production and for investigating the mechanics of tumor progression in vitro, Proceedings of the National Academy of Sciences 110(37) (2013) 14843–14848.

[25] R.L.F. Amaral, M. Miranda, P.D. Marcato, K. Swiech, Comparative Analysis of 3D Bladder Tumor Spheroids Obtained by Forced Floating and Hanging Drop Methods for Drug Screening, Frontiers in Physiology 8 (2017).

[26] C. Shao, J. Chi, H. Zhang, Q. Fan, Y. Zhao, F. Ye, Development of Cell Spheroids by Advanced Technologies, Advanced Materials Technologies 5(9) (2020) 2000183.

[27] C.S. Hughes, L.M. Postovit, G.A. Lajoie, Matrigel: A complex protein mixture required for optimal growth of cell culture, PROTEOMICS 10(9) (2010) 1886–1890.

[28] K. Stevenson, A.F. McVey, I.B.N. Clark, P.S. Swain, T. Pilizota, General calibration of microbial growth in microplate readers, Scientific Reports 6(1) (2016) 38828.

[29] M. Rebeaud, C. Bouche, S. Dauvillier, C. Attané, C. Arellano, C. Vaysse, F. Fallone, C. Muller, A novel 3D culture model for human primary mammary adipocytes to study their metabolic crosstalk with breast cancer in lean and obese conditions, Scientific Reports 13(1) (2023).

[30] M. Zanoni, F. Piccinini, C. Arienti, A. Zamagni, S. Santi, R. Polico, A. Bevilacqua, A. Tesei, 3D tumor spheroid models for in vitro therapeutic screening: a systematic approach to enhance the biological relevance of data obtained, Scientific Reports 6(1) (2016) 19103.

[31] B. Mittal, Subcutaneous adipose tissue &amp; visceral adipose tissue, Indian Journal of Medical Research 149(5) (2019) 571–573.

[32] K.L. Spalding, E. Arner, P.O. Westermark, S. Bernard, B.A. Buchholz, O. Bergmann, L. Blomqvist, J. Hoffstedt, E. Näslund, T. Britton, H. Concha, M. Hassan, M. Rydén, J. Frisén, P. Arner, Dynamics of fat cell turnover in humans, Nature 453(7196) (2008) 783–787.

[33] P. Arner, S. Bernard, M. Salehpour, G. Possnert, J. Liebl, P. Steier, B.A. Buchholz, M. Eriksson, E. Arner, H. Hauner, T. Skurk, M. Rydén, K.N. Frayn, K.L. Spalding, Dynamics of human adipose lipid turnover in health and metabolic disease, Nature 478(7367) (2011) 110–113.

[34] S.R. Farmer, Transcriptional control of adipocyte formation, Cell Metabolism 4(4) (2006) 263–273.

[35] E.D. Rosen, O.A. Macdougald, Adipocyte differentiation from the inside out, Nature Reviews Molecular Cell Biology 7(12) (2006) 885–896.

[36] M.I. Lefterova, M.A. Lazar, New developments in adipogenesis, Trends in Endocrinology & Metabolism 20(3) (2009) 107–114.

[37] A.G. Cristancho, M.A. Lazar, Forming functional fat: a growing understanding of adipocyte differentiation, Nature Reviews Molecular Cell Biology 12(11) (2011) 722–734.

[38] F. Baganha, R. Schipper, C.E. Hagberg, Towards better models for studying human adipocytes*in vitro*, Adipocyte 11(1) (2022) 413–419.

[39] V.M. Lauschke, C.E. Hagberg, Next-generation human adipose tissue culture methods, Current Opinion in Genetics &amp; Development 80 (2023) 102057.

[40] N. Shoham, A. Gefen, Mechanotransduction in adipocytes, Journal of Biomechanics 45(1) (2012) 1–8.

[41] N. Alkhouli, J. Mansfield, E. Green, J. Bell, B. Knight, N. Liversedge, J.C. Tham, R. Welbourn, A.C. Shore, K. Kos, C.P. Winlove, The mechanical properties of human adipose tissues and their relationships to the structure and composition of the extracellular matrix, American Journal of Physiology-Endocrinology and Metabolism 305(12) (2013) E1427–E1435.

[42] K. Duval, H. Grover, L.-H. Han, Y. Mou, A.F. Pegoraro, J. Fredberg, Z. Chen, Modeling Physiological Events in 2D vs. 3D Cell Culture, Physiology 32(4) (2017) 266–277.

[43] M.W. Laschke, M.D. Menger, Life is 3D: Boosting Spheroid Function for Tissue Engineering, Trends in Biotechnology 35(2) (2017) 133–144.

[44] E. Bellas, C.S. Chen, Forms, forces, and stem cell fate, Current Opinion in Cell Biology 31 (2014) 92–97.

[45] N.-E. Ryu, S.-H. Lee, H. Park, Spheroid Culture System Methods and Applications for Mesenchymal Stem Cells, Cells 8(12) (2019) 1620.

[46] C. Cero, W. Shu, A.L. Reese, D. Douglas, M. Maddox, A.P. Singh, S.L. Ali, A.R. Zhu, J.M. Katz, A.E. Pierce, K.T. Long, N. Nilubol, R.H. Cypess, J.L. Jacobs, F. Tian, A.M. Cypess, Standardized In Vitro Models of Human Adipose Tissue Reveal Metabolic Flexibility in Brown Adipocyte Thermogenesis, Endocrinology 164(12) (2023).

[47] I. Mora, F. Puiggròs, F. Serras, K. Gil-Cardoso, X. Escoté, Emerging models for studying adipose tissue metabolism, Biochemical Pharmacology 223 (2024) 116123.

[48] Y.-R. Liou, Y.-H. Wang, C.-Y. Lee, P.-C. Li, Buoyancy-Activated Cell Sorting Using Targeted Biotinylated Albumin Microbubbles, PLOS ONE 10(5) (2015) e0125036.

[49] C. Han, S. Takayama, J. Park, Formation and manipulation of cell spheroids using a density adjusted PEG/DEX aqueous two phase system, Scientific Reports 5(1) (2015) 11891.

